# Benchmarking the PEPOP methods for mimicking discontinuous epitopes

**DOI:** 10.1101/435974

**Authors:** Vincent Demolombe, Alexandre G. de Brevern, Franck Molina, Géraldine Lavigne, Claude Granier, Violaine Moreau

## Abstract

Computational methods provide approaches to identify epitopes in protein antigens to help characterizing potential biomarkers identified by high-throughput genomic or proteomic experiments. PEPOP version 1.0 was developed as an antigenic or immunogenic peptide prediction tool. We have now improved this tool by implementing 32 new methods (PEPOP version 2.0) to guide the choice of peptides that mimic discontinuous epitopes and thus potentially able to replace the cognate protein antigen in its interaction with an antibody. In the present work, we describe these new methods and the benchmarking of their performances.

Benchmarking was carried out by comparing the peptides predicted by the different methods and the corresponding epitopes determined by X-ray crystallography in a dataset of 75 antigen-antibody complexes. The Sensitivity (Se) and Positive Predictive Value (PPV) parameters were used to assess the performance of these methods. The results were compared to that of peptides obtained either by chance or by using the SUPERFICIAL tool, the only available comparable method.

The PEPOP methods were more efficient than, or as much as chance, and 33 of the 34 PEPOP methods performed better than SUPERFICIAL. Overall, “optimized” methods (tools that use the traveling salesman problem approach to design peptides) can predict peptides that best match true epitopes in most cases.

## Introduction

Antigen-antibody interactions are at the heart of the humoral immune response. B-cell epitopes correspond to the regions of the protein antigen that are recognized by the antibody paratope. Epitopes can be continuous (a linear fragment of the protein sequence) or discontinuous (constituted of several fragments scattered in the protein sequence, but nearby on the surface of the folded protein)[1–3]. Most protein epitopes are discontinuous [4,5] and the*refore*very difficult to map. Epitope identification and characterization are, however, pivotal steps in the development of immunodiagnostic tests [6], epitope-driven vaccines [7] and drug design as well as in protein function discovery, biochemical assays or proteomic studies for biomarker discovery. Epitopes can be mapped using various experimental methods [8–12] among which crystallographic analysis of antigen-antibody complexes is considered to give the most reliable information [13,14]. These techniques are, however, time-, resource- and labor-consuming, and, thus, unsuitable for proteomic applications. Computational methods could be an attractive alternative. B-cell epitope prediction methods [9,15–17] try to bioinformatically predict the antibody binding site on a protein sequence or on the 3D structure of a protein antigen. However, epitopes are not structural entities on their own. Epitopes and paratopes are relational entities that are defined by their mutual complementarity [18]. Thus, trying to predict *a priori* the identity of a protein epitope is a difficult task. For this reason, epitope predictors that take into account the sequence or structure of the antibody have been developed [19–21], but are of limited application since available antibody structures are scarce. Moreover, benchmark studies have highlighted that tools for predicting continuous epitopes have low efficiency [22–24] and that methods based on the antigen 3D structure show limited sensitivity (Se) and positive predictive value (PPV) [25].

We approached this issue from a slightly different point of view. Considering that the surface of a protein is a mosaic of potential antigenic epitopes, each of which could be bound by a cognate antibody [26], we developed the PEPOP 1.0 tool [27,28] to generate series of peptide sequences that can replace continuous or discontinuous epitopes in their interaction with their cognate antibody. Differently from discontinuous epitope predictors where the output prediction is either a list of amino acids (aa) or small protein fragments [29–31], PEPOP proposes peptide sequences that can be used directly in experiments. This tool promises to facilitate the manipulation of proteins in a way dealing with the output of proteomic studies.

We have previously validated the capacity of PEPOP 1.0 to generate immunogenic [27] and antigenic peptides that can be experimentally probed with antibodies to disclose the cognate epitopes [32–35].

As most antibodies against protein antigens recognize discontinuous epitopes, peptide design methods should take into account the structural information and try to guess (mimic) the epitope discontinuity. We thus improved the PEPOP tool (version 2.0) by focusing on methods for better predicting peptides aimed at mimicking discontinuous epitopes. It is now possible using PEPOP to generate large series of peptides that, collectively, should represent the accessible surface of the protein with its mosaic of putative epitopes. Consequently, within these large series of peptides, at least some should appropriately mimic antigenic epitopes.

In the present work, we describe these new methods and the benchmarking of their performances. To this aim, we used a comprehensive methodology and a series of test proteins for which epitopes have been experimentally determined by X-ray crystallography, which is the reference method. We show that the performance of each method is specific and that one method (TSPaa) performs better in these specific benchmarking conditions. We also compared the peptides designed by the different PEPOP methods with those predicted by SUPERFICIAL, in which the 3D structure of the protein surface is transformed into a peptide library [36], or by chance. PEPOP is available at http://pepop.sys2diag.cnrs.fr/.

## Results

### PEPOP principle

PEPOP is an algorithm dedicated to the design of peptides that are predicted to replace a protein epitope in its interaction with an antibody [27]. To bioinformatically design a peptide from the 3D structure of a given protein, a reference is chosen as a starting point. This can be a surface-accessible aa or a segment (i.e., a fragment of the protein composed of accessible and contiguous aa, from one to n aa, in the sequence) determined by PEPOP. After the identification of the aa or segments neighboring the reference, a method is used to delineate a path between them and to link them in order to generate the designed peptide. The aa or segments neighboring the reference are selected in an area of extension that can be either a cluster or a patch. To form a cluster PEPOP groups segments according to their spatial distances. A patch is defined around the reference. A requested peptide length has to be specified by the user in some methods. PEPOP proposes 35 methods: one method (the FPS method already included in PEPOP 1.0) generates peptides representing continuous epitopes, whereas the other 34 methods (of which NN and ONN were already present in PEPOP 1.0) are focused on peptides mimicking discontinuous epitopes.

### Methods’ redundancy

To assess the capacity of the different PEPOP methods to predict peptides that mimic epitopes, we used a dataset of experimentally (X-ray crystallography) determined epitopes that was filtered to eliminate any epitope redundancy (Table S1) [25]. The redundancy in the output sequences generated by the different PEPOP methods was verified by comparing the set of peptides predicted by each method (see “Peptide prediction” below). The low output redundancy by the different methods (Figure S2) indicated that the sequences of the generated peptides were highly diverse, except among methods of the same category. Peptides obtained using the SHP- and TSP-based methods showed less similarity with peptides obtained with the other methods (from 0% to 53% and 61%, respectively). The OPP, SHPaa and TSPaa methods were the most original methods because their peptides did not show any or only few similarities with the peptides generated by the other methods (37% at most). As the methods were developed to take into account different parameters, these results indicate that, except for few methods, sampling is large. The PEPOP methods are thus complementary, bringing diversity in the range of predicted peptides. This is useful when trying to represent the huge diversity of possible epitopes on a protein antigen.

### Peptide prediction

Each of the 34 methods was used to generate a series of peptide sequences from the 3D coordinates of the 75 antigens. As a protein is composed of a mosaic of epitopes [26,32], any region (cluster or patch) is potentially an epitope. Hence, all the possible peptides from a protein were predicted: each reference, segments or aa was used in turn to design a peptide. In this way, the whole protein surface was represented by the set of peptides generated by a given method.

### Requested length

To design a peptide using PEPOP, a sequence length for the predicted peptide has to be chosen (requested length). Nonetheless, as segments of variable lengths are used to build the peptide sequence, the final peptide length might differ from the requested length. The appropriate requested length to use for benchmarking was determined by requesting discrete lengths (ranging from 8 to 16 aa) for the predicted peptides. The final mean peptide sizes are reported in Table 1. As expected, the average final lengths were higher than the requested lengths by 2 or 3 aa and increased with the requested length. According to the chosen method, the average final peptide lengths could be very different. For benchmarking, the prime and linker methods should use the same requested length, unless this lead to different peptide sizes, to allow their comparison and the evaluation of the linker contribution. For the evaluation, two requested lengths were chosen: 12 aa for the prime and linker methods and 16 for the graph-based methods because they lead to an average final peptide length close to the mean length of the epitopes in the dataset (16.7 aa).

**Table 1.** **Mean** peptide size according to the requested peptide length.

| mean | L=8 | L=10 | L=12 | L=14 | L=16 |
| --- | --- | --- | --- | --- | --- |
| of the means | 11.66 | 13.35 | 14.95 | 16.45 | 17.70 |
| standard deviation | 2.16 | 2.34 | 2.66 | 3.02 | 3.29 |
| Prime methods | 10.44 | 11.97 | 13.53 | 15.14 | 16.65 |
| ALA methods | 14.98 | 17.24 | 19.06 | 20.36 | 21.05 |
| SA methods | 11.45 | 13.13 | 14.77 | 16.44 | 17.88 |
| SAS methods | 12.96 | 14.87 | 16.72 | 18.47 | 19.80 |
| Prime and Linker methods | 12.48 | 14.33 | 16.05 | 17.62 | 18.84 |
| Graph-based methods | 10.16 | 11.55 | 12.96 | 14.31 | 15.59 |
| SHP based methods | 13.27 | 13.27 | 13.27 | 13.27 | 13.27 |
| TSP based methods | 9.13 | 10.98 | 12.85 | 14.66 | 16.37 |
| TSPaa method | 8.00 | 9.99 | 11.99 | 13.98 | 15.97 |
| TSPnat and TSPrev methods | 9.27 | 11.10 | 12.96 | 14.75 | 16.42 |

### Benchmarking

The 34 methods predicted a total of 119277 peptides (i.e., about 3508 per method and 1590 per antigen), using the 75 protein antigens of the dataset (Table S1). The Se (proportion of peptide residues present also in the epitope) and the PPV (proportion of epitope residues in the predicted peptide) parameters were used to evaluate the methods’ performance. A perfectly accurate prediction would give Se and PPV values of 1. The mean Se ranged from 0.34 to 0.49, according to the method, and the mean PPV was a little higher (between 0.39 and 0.57) (Figure S4). In the evaluation of discontinuous epitope prediction tools carried out by Ponomarenko & Bourne using the same dataset, the mean Se and PPV with the best method were 0.46 and 0.40, respectively [25], indicating that many of the PEPOP approaches are more efficient.

The Se and PPV mean values give a measure of the adequacy between the predicted peptide and the reference epitope, but they do not discriminate between methods. However, a researcher would wish to have a method that provides the highest possible number of peptides for the highest possible number of epitopes. To know whether a method is more performing than another, (i.e. whether a given method can predict the best possible matching peptides for the widest possible range of epitopes), the Se and PPV minimal value (threshold) to consider a method as theoretically efficient must be determined.

To select an appropriate threshold, we studied the distribution of peptides relative to their Se and PPV (Figure 2 and Figure S5). On average, a method predicted 7.6% of peptides with a Se and PPV above 0.6, 1.73% of peptides with a Se and PPV above 0.7 and 0.13% of peptides with a Se and PPV above 0.8 (i.e., about 267, 60 and 4 peptides, respectively, based on the mean number of peptides predicted per method). We finally selected the 0.7 value as the threshold because it offers a good compromise between “quality” and quantity of predicted peptides.

**Figure 1.**
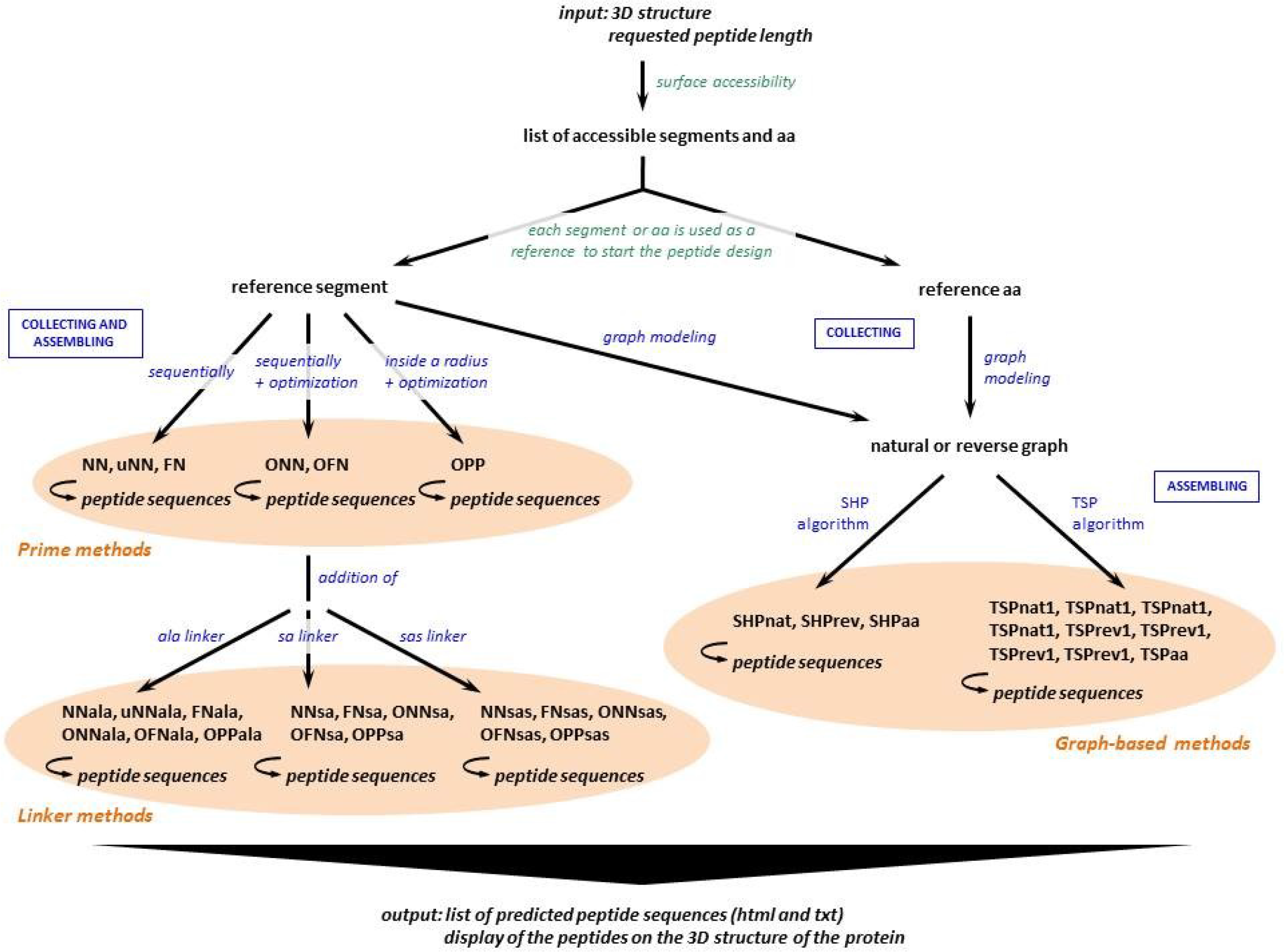
Flowchart describing how PEPOP predicts a series of peptide sequences (“Peptide Bank” section of the web site of PEPOP).

**Figure 2.**
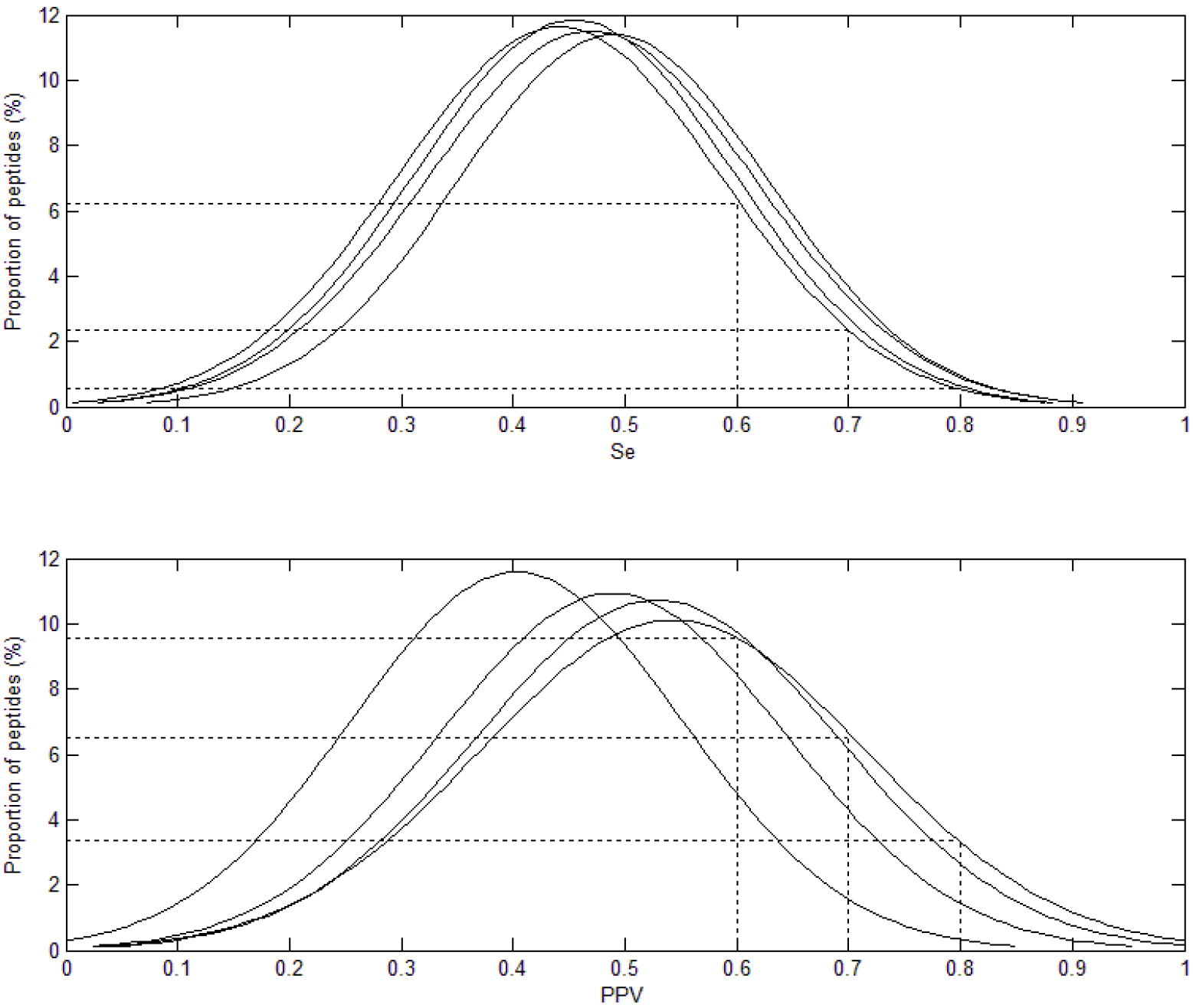
Example of the distribution of the Se (upper panel) and PPV (lower panel) values of the peptides predicted by the OFN methods (OFN, OFNala, OFNsa, OFNsas) without taking into account the aa positions.

We then calculated the proportion of peptides with a Se and a PPV higher than 0.7 for each method (Figure 3, empty bars). Group of methods were roughly clustered around similar values. Indeed, 1.4 to 1.9% of peptides generated by using the prime methods had Se and PPV values above 0.7, whereas this percentage decreased to 0.7% for peptides designed with the ALA methods. Compared to the prime methods, the performance of the ALA methods decreased due to the beneficial effect of the addition of Alanine residues between segments on Se and its unfavorable effect on PPV. The SA methods were slightly more efficient than the prime methods, but not the SAS methods. This also was the result of a beneficial effect of the addition of aa linkers on Se and their negative effect on PPV. TSP-based methods, particularly TSPaa, were the most efficient as they generated the highest percentage of peptides with Se and PPV above 0.7.

**Figure 3.**
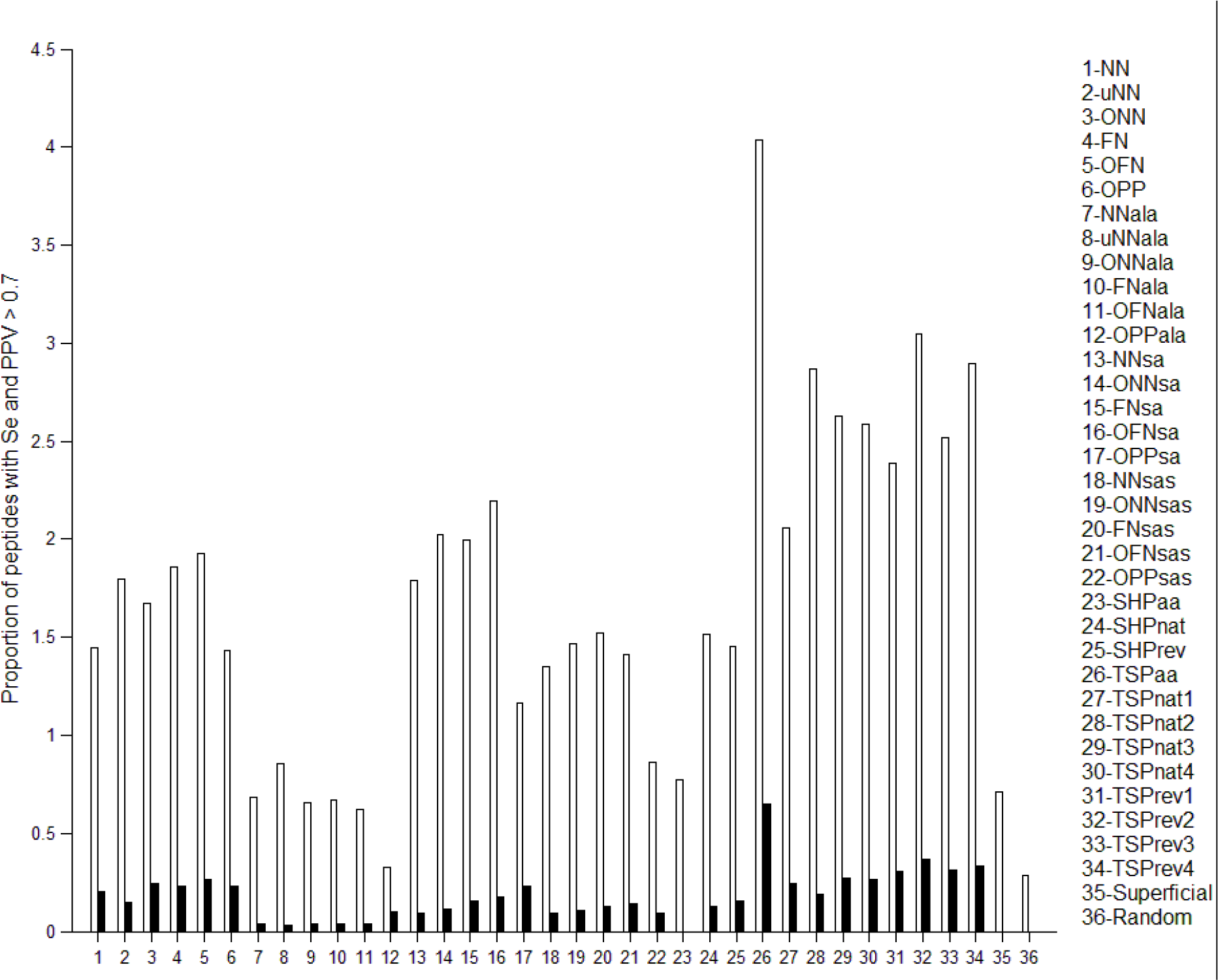
Performances of the methods: proportion of peptides with Se and PPV > 0.7. Empty bars, the aa positions were not taken into account; solid bars, the aa positions were taken into account.

We then calculated for how many antigens, a given method would generate peptides with a Se and a PPV higher than 0.7 (Fig. 4, empty circles) in order to know whether a method was efficient with different proteins. As before (see Fig. 3), methods from the same group showed similar performances and the most efficient were the TSP methods. Indeed, TSPaa, TSPnat3 and TSPrev4 targeted the highest number of antigens.

**Figure 4.**
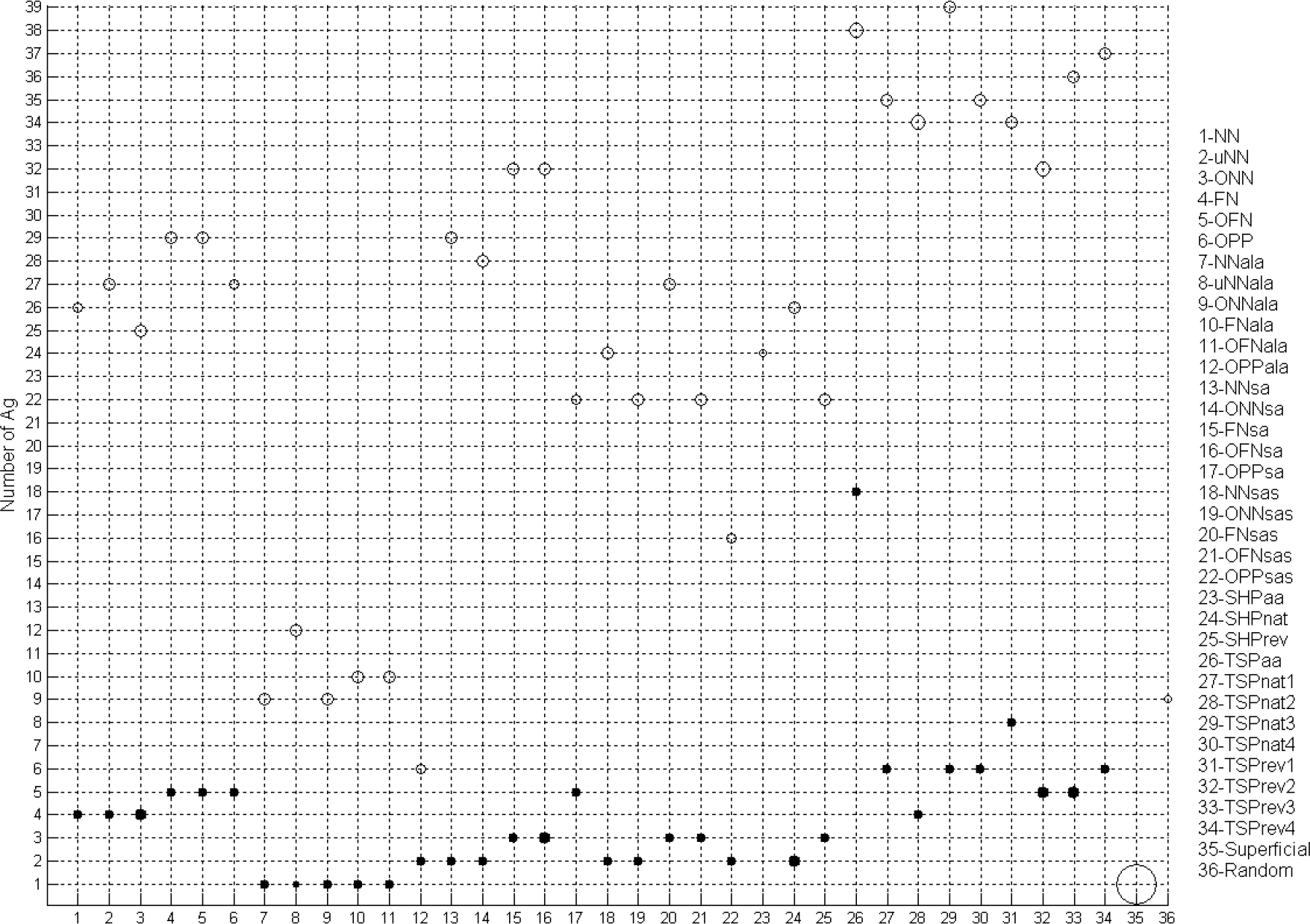
Robustness of the performance of the methods. For each method, the number of antigens is plotted with a circle size proportional to the number of peptides having Se>0.7 and PPV>0.7. Empty circles, the aa positions have not been taken into account; solid circles, the aa positions have been taken into account.

#### Influence of peptide length

As Wang and collaborators [37] showed that their performance classification was dependent on the epitope length, we studied the influence of the peptide length on the methods’ performance. First, we determined the number of peptides with Se and PPV above 0.7, relative to the peptide-epitope size difference (Fig. 5). We found that peptides that were closer in size to the epitope performed better. We then analyzed the influence of the five requested peptide lengths (8, 10, 12, 14 and 16 aa) on the performance of the methods (Fig. 6). The peptide length had, as expected, no influence on the performance of the SHP and OPP methods (because the final peptides are identical whatever the requested peptide length) (Figure 6C). It had a weak influence on the performance of the ALA methods (the final peptides are longer than the requested length, but they are only enriched in Ala residues) and on the SA and SAS methods (peptides are possibly enriched of several aa, and this may have an unfavorable effect depending on the epitope composition). Conversely, the performance of the NN, NNu, ONN, FN, OFN and TSP methods progressively increased with the peptide length. Nevertheless, TSPaa remained the most performing method. These results also show that when selecting the PEPOP parameters, it is advisable to request a peptide length close to the epitope size, i.e. a number of aa of the peptide close to the average number of aa contained in an epitope (17aa).

**Figure 5.**
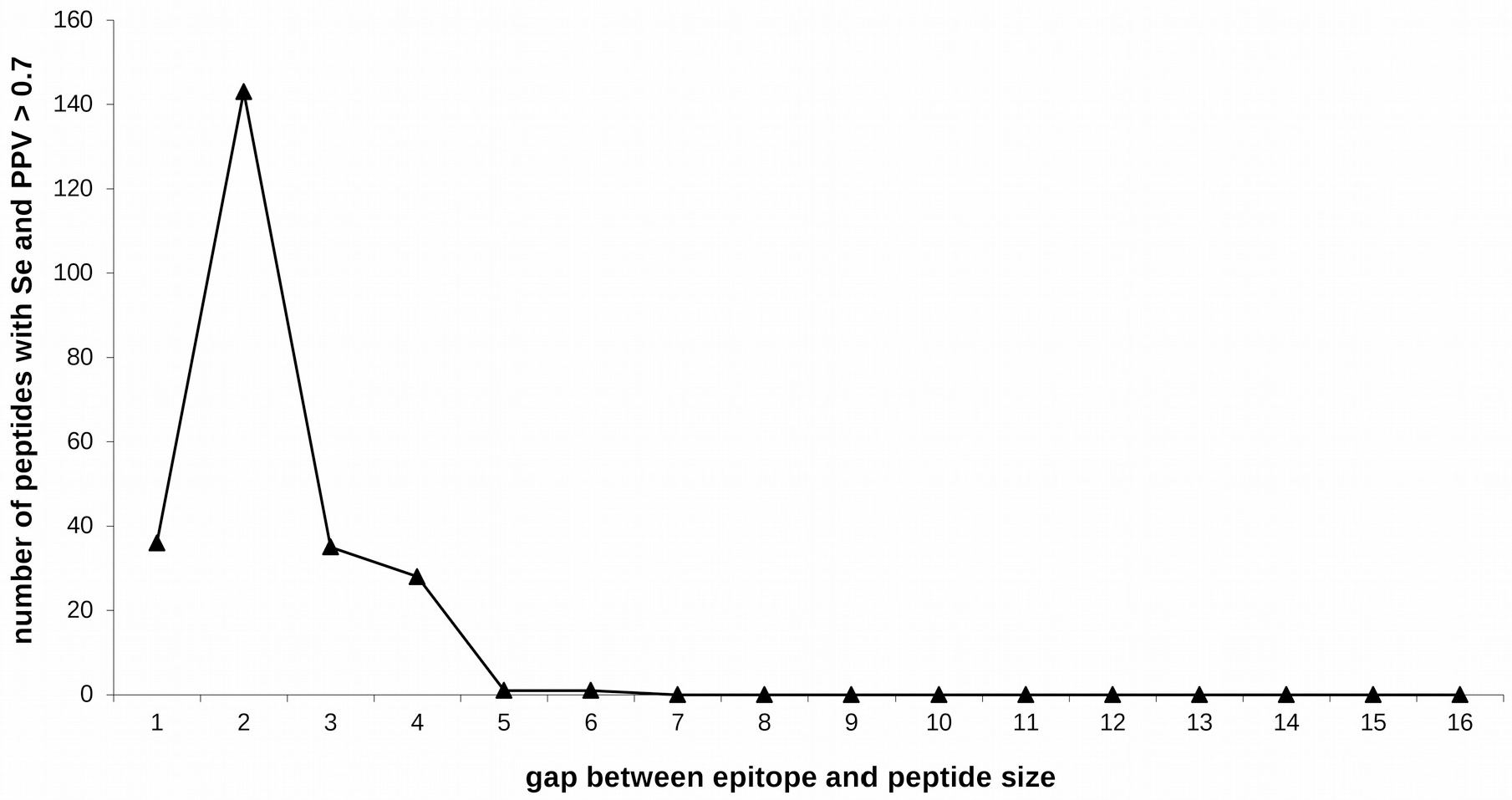
Relationship between peptide performance and size similarity between epitope and peptide. Aa positions were taken into account.

**Figure 6.**
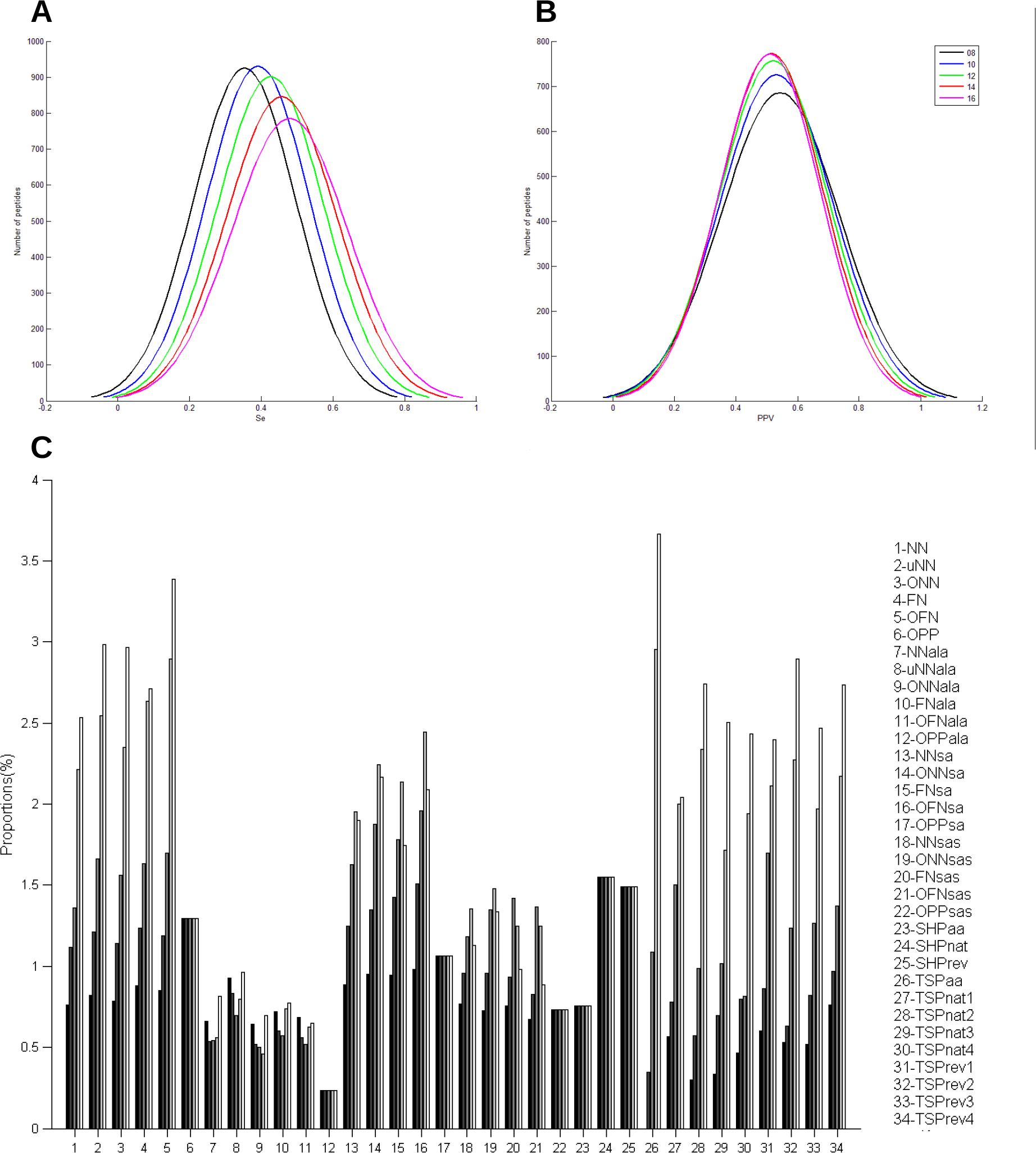
Influence of the requested peptide length on the methods’ performance. **A.** Se and **B.** PPV distribution according to the requested peptide length. **C.** Proportion of peptides with Se and PPV >0.7 based on the requested peptide length, from 8 (solid bars) to 16 (empty bars) aa with an increment of 2 at each step.

#### Amino acid positions

The aa positions were not taken into account for Se and PPV computations because the nature of the aa and not their original position in the protein is important for antibody recognition. However, the closeness and the order of aa residues in the peptide could be important factors for protein mimicry by peptides. Thus, we computed again the Se and PPV values by taking into account the aa position in the predicted peptides compared to their position in the epitope. As expected, the Se and PPV values of all methods decreased when taking into account the aa positions (Fig. 3, black bars, and Fig. 4, black circles, and Figure S4). All methods showed comparable mean PPV values, except for the SA and SAS methods (Figure S4A). The mean Se values, when taking or not into account the aa positions, have similar profiles (Figure S4B). Despite the overall reduction in efficiency (i.e., proportion of peptides with both Se and PPV higher than 0.7) when taking into account the aa positions, methods followed the same tendency as the analysis that did not take into account the aa positions. TSPaa again was the most efficient method (Fig.3). When calculating for how many antigens a given method would generate peptides with a Se and a PPV higher than 0.7 by taking into account the aa positions, the most efficient method was TSPrev1 instead of TSPnat3, but overall, the TSP methods still performed best (Fig.4). On the other hand, SHPaa did not produce any efficient peptide (Fig.3 and 4). Thus, the results of the Se and PPV computations that take into account the aa positions confirmed the previous analysis.

#### Comparison with SUPERFICIAL

We then compared our results with those obtained using SUPERFICIAL (Sfs), the only other available peptide design tool. This software predicted 143 discontinuous peptides from 30 antigens of the dataset, including 21 peptides from only one antigen. The proportion of peptides generated by SUPERFICIAL with Se and PPV values higher than 0.7 was about 0.7% (Figure 3), a value similar to the one obtained with the lowest performing PEPOP methods. Moreover, SUPERFICIAL did generate peptides with Se and PPV values higher than 0.7 only for one antigen among the 75 proteins of the dataset (Fig. 4). Finally, SUPERFICIAL did not predict any peptide with Se and PPV higher than 0.7 when the aa positions were taken into account (Fig.3). In conclusion, all PEPOP methods performed better than the only other available peptide design tool.

#### Comparison with chance

Finally, we compared the PEPOP methods to a method that predicts peptides by chance. The performance of the random method was comparable to that of the less efficient PEPOP methods (ALA methods). It predicted 0.8% of peptides with Se and PPV values higher than 0.7 for one antigen out of 3. When the aa positions were taken into account, the random method did not predict any efficient peptide (Fig.3). These results show that peptides predicted by the PEPOP methods are not efficient only by chance. This was further confirmed by the probability to predict the most efficient peptide (10E-24). These results indicate that the PEPOP methods perform much better than chance.

### Example

We then wanted to assess how well the most efficient peptides (best Se and PPV) designed by PEPOP represent the corresponding epitope on the 3D structure of its antigen (Figure 7). Among all the PEPOP methods, we selected the peptide having the best Se and PPV without taking into account the aa positions (Fig.7A) and the peptide having the best Se and PPV when taking into account the aa positions (Fig.7B). The two peptides matched pretty well their epitope (all the epitopic aa are predicted in the peptides). Nonetheless, the Se and PPV values greatly changed depending on how they were calculated. For example, the first peptide has Se and PPV of respectively 1 and 0.864 but if the aa positions are not taken into account, Se and PPV are of only 0.368 and 0.318 respectively. However, the peptide includes effectively the same nature of aa than the epitope. This example shows that what is important for the final peptide is the nature of the aa, not its position in the protein.

**Figure 7.**
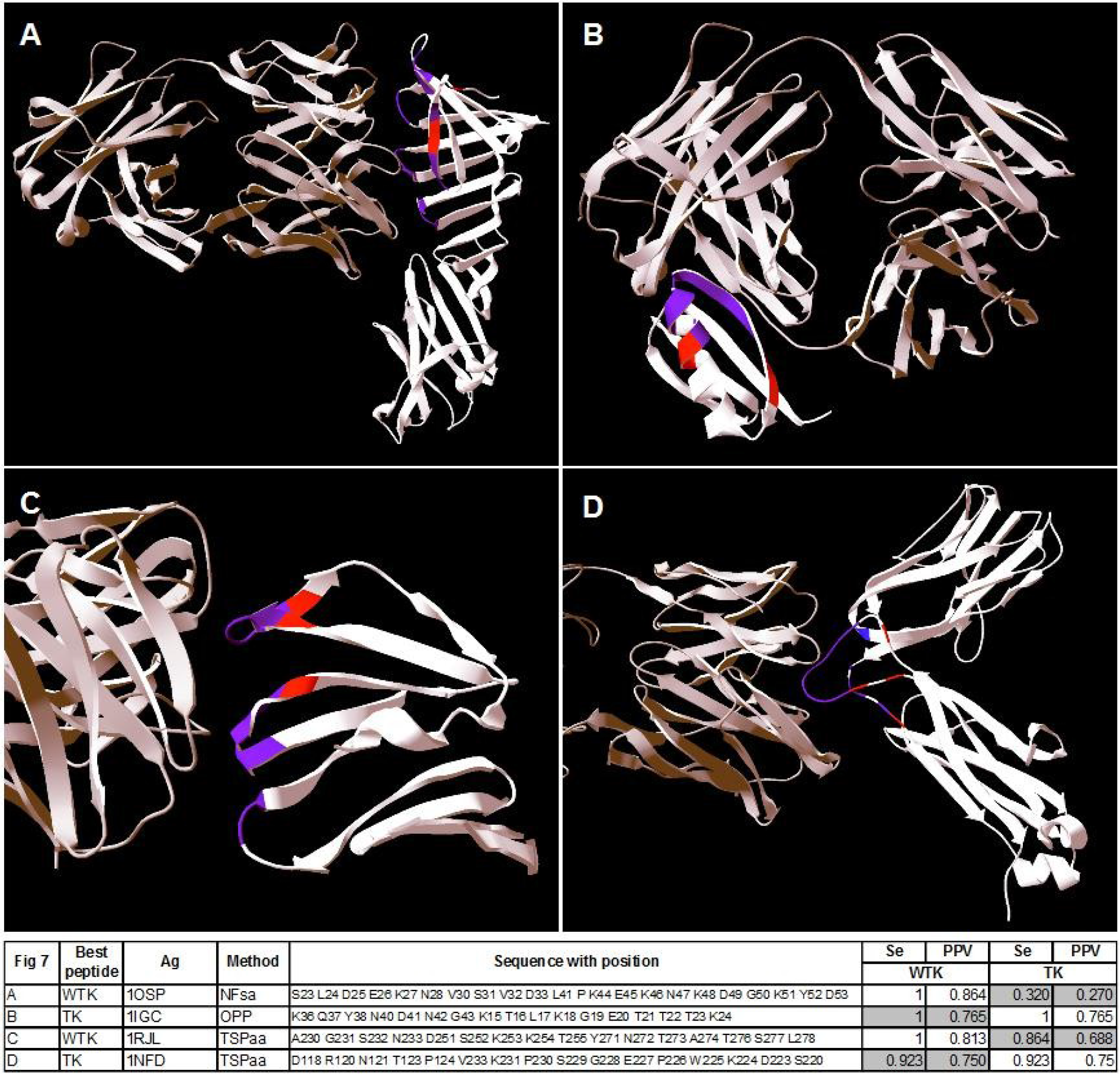
3D views of the most efficient peptides generated with the dataset using all PEPOP methods (**A** and **B)** or TSPaa (**C** and **D**). A and C, peptides having the best Se and PPV computed without taking into account the aa positions (WTK); B and D, peptides having the best Se and PPV computed by taking into account the aa positions (TK). The peptide aa are in red, the epitope aa are in blue and common aa are in purple. The antibody is in grey.

We then selected the two best peptides (highest Se and PPV) generated by TSPaa method when taking (Fig. 7D) or not (Fig.7C) into account the aa positions. These two predicted peptides also match pretty well their epitope (only one epitopic aa is not predicted in the peptides). This example shows that even if TSPaa is the best performing method, it does not actually predict, for every antigen, the most efficient peptides since these two peptides have Se and PPV slightly lower than the two previous peptides. Hence, all the PEPOP methods should lead to efficient peptides. In any case, the peptides contained a majority of epitopic aa and thus they should be recognized by the antibody, particularly because the arrangement between aa was optimized.

## Discussion

PEPOP (http://pepop.sys2diag.cnrs.fr/) is a prediction tool of antigenic / immunogenic peptides. It was developed, not to predict epitopes, but to deliver series of peptides that should mimic epitopes, particularly discontinuous epitopes, which are the more common ones. It is more complex to predict peptides mimicking discontinuous than continuous epitopes because the aa order is not already defined. As enumerating all possible peptides would amount to solving a complex NP-complete problem and it would anyhow be impossible to test all of them computationally or experimentally, a limited enumeration must be defined. To this aim, PEPOP version 2.0 was improved by implementing 32 new methodologies that exploit different criteria, such as the distance between segments, their disposition in the peptide and their conformation relative to the protein antigen, to design discontinuous peptides that match as much as possible the antigen. The main principle is to find a path, an arrangement, between elements (segments or aa) of a defined area on the protein that will compose the final peptide.

The prime methods design a peptide from a reference segment and add to it its neighboring segments. Compared to the NN method, the FN method was developed to maintain the reference segment in the central position. The ONN, OFN and OPP methods search for the most natural path between segments by minimizing the traveled distance. The linker methods add (or not) aa between segments. The purpose of the ALA method is to keep in the peptide the same segment spacing of the antigen protein to allow the interacting aa of the antibody to establish contacts. The SA and SAS methods use the protein blocks (PBs) of a structural alphabet [38,39] to facilitate the adoption of the protein conformation. To bypass the NP-complete problem of enumerating all possible arrangements between the segments composing the peptide (n! permutations) in ONN, ONF and OPP methods, we used the graph theory. In the graph-based methods, the objective is to find the optimal path between segments or aa, as this should lead to peptides close to the native protein context.

Here, we evaluated the performance of the 34 methods included in PEPOP 2.0 by measuring the match between the peptide composition and that of known discontinuous epitopes and classified as efficient the methods that predicted the largest number of peptides with both Se and PPV values higher than 0.7 (efficient peptides). We found that the TSP-based methods, particularly TSPaa, TSPnat3 and TSPrev2, predicted the best matching peptides in most cases, although they did not lead to the best peptide (Figure 7). TSPaa was the most efficient method *in silico* as it predicted the largest percentage of efficient peptides for the highest number of antigens. These methods performed better because the search of the optimal path using the TSP allows selecting the correct segment or aa (i.e., the segment or aa present in the epitope). All PEPOP methods, except OPPala, were more successful than SUPERFICIAL and more efficient than or as much as the method predicting peptides by chance.

Benchmarking of different computational methods must be done with precaution as the tools, datasets and metrics can be different from one analysis to the other, thus not allowing objective comparisons. Even the definition of epitope can be different. Indeed, some consider the part of the antigen recognized by one antibody as an epitope on its own, whereas others consider to be an entire epitope all the aa found to interact with any antibody [16]. Moreover, some authors think that proteins have only one or few epitopes on the surface [40], whereas others see a protein as a mosaic of epitopes [26]. Finally, because all the possible epitopes could not be discovered, only a few of the features that characterize epitopes are used when trying to discriminate between epitopic and non-epitopic aa. This could at least partially explain why all epitope prediction tools show a weak performance [22,25]. And, we believe that an epitope cannot be faithfully predicted without taking into account its antibody partner, because epitopes only exist through the interaction with their cognate antibody [41]. Thus, studies taking into account the antibody partner in predicting antigen epitopes are of particular interest [19–21,42] although, because of the poor availability of antibody data, they currently cannot be applied in high-throughput analyses.

To determine whether some of the 34 methods included in PEPOP version 2.0 for designing discontinuous peptides are more relevant than others, we carried out a benchmark process. To this aim we followed as much as possible the recommendations by Greenbaum and collaborators [43] for assessing the performances of epitope prediction methods, although PEPOP goal is slightly different from that of “classical” epitope prediction tools. We decided to use Se and PPV together to select the most efficient peptides, although they are threshold-dependent. Indeed, when used on their own, they do not provide a complete picture of the method performance. For instance, a peptide with a Se (number of epitopic aa included in the peptide) close to 1 could also contain many additional aa that might disturb its recognition by the antibody. Similarly, a peptide with a PPV (number of the peptide aa included in the epitope) close to 1 could contain not enough epitopic aa for antibody recognition. Considering a given antigen-antibody interaction, not all generated peptides will match the epitope because peptides come from the entire surface of the protein. However, a method can be considered efficient if it yields an elevated number of peptides that closely match the epitope (i.e., with both Se and PPV higher than 0.7 in our study).

Nevertheless, benchmarking under-evaluated the linker-based methods. Indeed, even if a peptide generated using these methods included all the aa of the epitope, its PPV would be lower than the PPV of the same peptide without linker (e.g., a peptide designed using the ONNala, ONNsa or ONNsas method versus the same peptide generated using ONN). This despite the fact that the linker methods were developed to increase the performance, based on the hypothesis that spacing the segments by linker amino acids would better mimic their real disposition on the protein and consequently facilitate the peptide recognition by the antibody. Although this bias was compensated by a slightly higher Se, we feel that the principle on which these methods are based has been not perfectly appraised.

It is difficult to claim that one method is better than another one. Indeed, the good performances of one method in terms of Se and PPV do not ensure that the corresponding peptides will actually be recognized by an antibody. Only their experimental evaluation can confirm the peptide reactivity. Indeed, the idea behind the PEPOP tool is that, due to the inherent difficulty to guess an epitope, it would be preferable to generate a comprehensive series of peptides that can be experimentally assessed to determine which ones are endowed with the properties of a functional epitope. The mean number of peptides predicted per antigen by the PEPOP methods was 1590. Experimentally testing the antigenicity of about 1500 peptides is feasible by techniques like peptide microarrays [44–46]. Thus, the specific epitopes of a given antigen could be identified by running all PEPOP methods, synthesizing the generated peptides and testing them in microarrays. Conversely, the experimental validation of about 120000 peptides (number of peptides designed by the PEPOP methods for the entire dataset) would require too much time and resources and probably would not be feasible for any future peptide design tool.

Testing the immunogenicity of at least 1500 peptides would be even worse. Due to these difficulties to realize systematic experimental validations, we believe PEPOP, and others similar tools, have to be seen as “test tubes” which will gradually be validated as studies will be developed, until a consensus satisfactory validation process is developed. Although some studies begin to explore this problem [47] the proposed benchmark is not applicable for all epitope prediction tools neither for all studies. Anyway, PEPOP has already been successfully used in several studies of different goals [27,32–35,48,49].

In the workshop report by Greenbaum et al., Dr Van Regenmortel “emphasized the need to clarify the purpose of making a specific epitope prediction, and how this clarification could direct selection of the most appropriate prediction tool or development of a new tool, as needed”. PEPOP has been or can be used for all the purposes where surrogate epitopes are needed, purposes such as those cited by Van Regenmortel, i.e. “seeking vaccine candidates” [49] or “replacing antigens in diagnostic immunoassays”. It can also efficiently help in mapping epitopes [33–35,50,51] and would be a very informative tool for understanding the rules of molecular mimicry, a very difficult [23,41,52] but promising research field as testified by the number of available studies [9,53–56] and tools [30,57,58]. PEPOP could also help characterizing all new proteins discovered by high-throughput technologies, such as proteomics [59,60], by facilitating their manipulation.

## Experimental procedures

### Structural data

The dataset of 165 X-ray-determined epitopes was from (Ponomarenko & Bourne, 2007). To avoid bias caused by the over- or under-representation of an epitope described by several 3D structures of the same antigen-antibody complex, epitope redundancy was eliminated by keeping only one crystallography of a given antigen-antibody complex. Therefore, 90 antigen-antibody complexes were rejected. The final dataset was of 75 unique antigen-antibody complexes (Table S1).

The epitope size varied from 4 to 23 residues with only one exception (52 aa). The average size of an epitope was 16.7 aa (median: 17 aa). Epitopes were all discontinuous and were composed of 3 to 14 segments, each containing 1 to 12 contiguous aa. An epitope contained on average 7 to 8 segments of 2.38 aa. These data are in accordance with the literature [56,61,62].

### Epitope definition

An epitope was defined as a series of aa included in the protein antigen. These aa contained at least one atom that establishes a contact (i.e., a distance threshold lower than or equal to 4Å) with an atom from the antibody.

### PEPOP methods

Peptides that mimic the discontinuous epitopes of the dataset were designed using the different PEPOP methods (Figure 1). To build a peptide, the PEPOP algorithm concatenates either segment sequences (a continuous stretch of surface-accessible aa) or single surface-accessible aa from the antigen 3D structure. Based on Euclidian distances, the PEPOP methods first select the neighboring segments or aa and then determine in which order assemble them to form the final linear peptide sequence supposed to mimic the discontinuous epitope. PEPOP 2.0 has been improved by addition of 34 new methods to the two already present in PEPOP 1.0. These methods are based on different criteria because precise rules for peptide design are lacking due to our poor understanding of the mechanisms underlying the molecular mimicry of a native protein by a linear peptide. These methods can be classified in three main groups: (a) prime methods, (b) linker methods and (c) graph-based methods.

a. In the **prime methods** (Figure 1 and Figure S1) neighboring segments are collected around a reference segment (starting segment). Therefore, starting from the reference segment and until a defined peptide length is reached, the methods concatenate the segments as follows:

- **the nearest neighbor (NN) method** adds the sequence of the nearest neighbor segment C-terminally to the forming peptide;
- **the upset nearest neighbor (uNN) method** adds the sequence of the nearest neighbor segment C-terminally in the natural or the reverse sense according to the distance of the C-terminus of the forming peptide;
- **the flanking nearest neighbor (FN) method** adds the sequence of the nearest neighbor segment in turn C-terminally and N-terminally to build the peptide. More sophisticated prime methods are directly derived from these three firsts methods and are used to determine the optimized path between segments, i.e. in which order assemble them, by enumerating all possible arrangements. The sums of the distances between segments are then calculated. The optimized path corresponds to the arrangement with the shortest total distance. No extra aa is added. Thus, the optimized path can be calculated by using:

- **the optimized nearest neighbor (ONN) method** for segments found using the NN method.
- **the optimized flanking nearest neighbor (OFN) method**: for segments found with the FN method.
- **the optimized patched segments path (OPP) method:** for the set of segments present in a 10Å-radius patch.
b. **Linker methods.** As in the prime methods no intermediate aa is added, 16 **linker methods** were then derived from these methods to add extra aa (Figure 1 and Figure S1) between segments generated by one of the prime methods (NN, ONN, FN, OFN, OPP):

- **the ALA linker methods** (NNala, uNNala, ONNala, FNala, OFNala, OPPala) add an alanine linker, as many times as the distance between segments allows the insertion of a peptide bond. Alanine is often considered as the most average aa in terms of length, volume and polarity.
- **the structural alphabet-based linker (SA) methods** (NNsa, ONNsa, FNsa, OFNsa, OPPsa) add zero, one or two aa as linkers. Protein blocks (PBs) [38,39] form a library of 16 small protein fragments of five residues in length that can approximate every part of a protein structure. PBs are overlapping, so each PB is followed by a limited number of PBs (i.e., some specific transitions exist between PBs). First, the transitions between segments are verified in the segments transcribed into PBs, based on the protein 3D structure. According to the PB transition matrix [63], if the transition between the last PB of a segment and the first PB of the following segment is allowed, no aa is added between these segments. If the transition is not allowed, a PB is virtually added by searching the one leading to the best PB transition. Adding a PB means adding an aa. The most favorable aa is determined from data calculated from the PDB file that reports each aa statistical preferences for each positions in each PB [38]. If a transition cannot be found, the process is repeated by adding two PBs. If also this does not work, the peptide is not possible.
- **the structural alphabet superposition-based linker (SAS) methods** (NNsas, ONNsas, FNsas, OFNsas, OPPsas) add one aa as linker according to the structural superposition of the segment using the structural alphabet approach to facilitate the peptide folding in the same fold as the corresponding fragments in the protein.
c. **Graph-based methods**(Figure 1) use the graph theory to model a given protein, its segments and its aa, in order to find the neighboring segments or aa. They employ three different graphs where edges are weighted by Euclidian distances. The first graph (“natural” graph) is oriented. The nodes are protein segments and can only be added in their natural sense, from N-terminus to C-terminus. The second graph (“reversed” graph) is the non-oriented version of the previous one. The third graph (“aa” graph) is non-oriented and the nodes are surface-accessible aa instead of segments. Two algorithms used these graph to find the optimal path, i.e. in which order assemble the segments or aa:

- **SHortest Path (SHP)-based methods** (three methods): from a set (i.e., a cluster or a patch, see definitions below) of elements (segments or aa), a peptide is the shortest path between two elements that include most aa residues. The **SHPnat** method uses the “natural” graph, **SHPrev** the “reversed” graph and **SHPaa** the “aa” graph.
- **Traveling Salesman Problem (TSP) based methods** (nine methods): from a set (cluster or patch) of elements (segments or aa), the TSP algorithm is used to find the optimal path (shortest distance) between elements. The **TSPnat** methods use the “natural” graph, the **TSPrev** methods use the “reversed” graph and the **TSPaa** method uses the “aa” graph. From the optimal path between the elements defined by the TSP algorithm, all possible peptides of the requested length are computed. The final peptide, identified using the:

▪ **TSPnat1** and **TSPrev1** methods, is the peptide with the highest score (see “peptide scoring”)
▪ **TSPnat2** and **TSPrev2** methods, is the peptide with the shortest traveled distance
▪ **TSPnat3** and **TSPrev3** methods, is the peptide for which the traveled distance according to the number of segments of the peptide is the shortest
▪ **TSPnat4** and **TSPrev4** methods, is the peptide that includes the two closest segments
▪ **TSPaa** method, is the peptide for which the traveled distance is the shortest.

The protein area used in the SHPnat, SHPrev, TSPnat and TSPrev methods is a cluster or a 15Å-radius patch (see definitions below). The area used in the SHPaa and TSPaa methods is a varying patch.

### Definition of cluster and patch

In PEPOP, clusters are segments grouped according to their spatial distances. They are calculated using Kitsch from the PHYLIP package v3.67 [64], as previously described [27]. PEPOP uses three types of patches. The 10Å- and 15Å-radius patches gather segments within a fixed distance, respectively 10Å and 15Å, from the center of gravity of a reference segment. The third patch type gathers the aa at a distance that varies from 15 to 20Å from a reference aa: the final radius is the one in which the average number of aa between radius 15, 16, 17, 18, 19 and 20Å is collected.

As each segment is used in turn to define a patch, the number of 10Å- and 15Å-radius patches is equal to the number of segments (Figure S3). The number of varying patches is equal to the number of accessible aa because each aa is used in turn to define a varying patch.

### Peptide scoring

In PEPOP, the score of a peptide is the sum of the scores of the segments composing the peptide [27]:

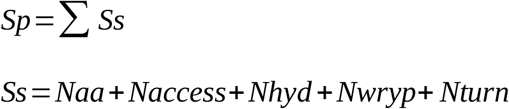

where Sp is the peptide score, Sp the segment score, Naa the number of amino acids composing the segment, Naccess the average accessibility of the segment, Nhyp the number of hydrophobic amino acids, Nwryp the number of specific amino acids (W, R, Y or P) and Nturn the number of amino acids involved in a β-turn.

### Peptide predictions

Depending on the PEPOP method, each segment or surface-accessible aa of a protein antigen is used as a reference to design a peptide.

In the **prime** (NN, uNN, ONN, FN, OFN, OPP methods) and **linker methods** (ALA, SA and SAS methods), peptides are predicted from each segment defined by PEPOP. The number of peptides generated by these methods corresponds to the number of segments.

In the **graph-based methods** that model protein segments (SHPnat, SHPrev, TSPnat1, TSPnat2, TSPnat3, TSPnat4, TSPrev1, TSPrev2, TSPrev3 and TSPrev4), peptides are predicted in PEPOP clusters and in 15Å-radius patches. As each protein segment is successively considered as a reference segment, there are as many patches as segments. The number of peptides predicted by these methods corresponds to the number of clusters plus the number of segments.

In the graph-based methods that model the surface-accessible aa of the protein (SHPaa and TSPaa), peptides are predicted in PEPOP clusters and in varying patches. The number of aa is computed for all radii between 15 and 20Å (1Å increment per step), and the radius leading to the average number of aa defines the final patch. The number of peptides predicted by these methods corresponds to the number of clusters plus the number of segments.

In each method, redundant peptide sequences are eliminated; however, two different methods can predict the same peptide sequence.

### Performance evaluation metrics

The capacity of each peptide generated by a given PEPOP method to mimic the epitope described in the reference dataset for that protein was evaluated using two criteria: the sensitivity (Se) and the positive predictive value (PPV).

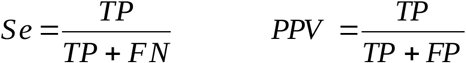

TP (True Positive) is the number of aa belonging to both the generated peptide and the reference epitope. FP (False Positive) is the number of aa belonging to the peptide, but not to the epitope. FN (False Negative) is the number of aa belonging to the epitope, but not to the peptide. Hence, Se represents the proportion of epitope aa present in the peptide, whereas PPV is the proportion of peptide aa present in the epitope.

The aa nature or the aa position in the peptide and reference epitope was then compared by not taking and by taking into account the aa positions of the protein. The aa used in linker methods (ALA, SA and SAS) were considered in the evaluation that takes into account the aa positions only after all the other aa of the peptide were compared with the epitopic aa. We chose to take into account the supplementary aa, because otherwise it would have amounted to evaluate again the results of the prime methods. Indeed, the only difference between prime and linker methods is the aa that are added between segments and that do not correspond to any position in the protein.

To measure the correlation between performance and size between peptides and epitopes, the absolute value was calculated.

### Chance

#### Random method

Peptides were designed as sequences of randomly selected aa according to the protein aa composition. The peptide length was randomly computed according to the distribution of peptide lengths designed by PEPOP using the dataset of 75 antigens. The number of peptides was randomly chosen according to the number of peptides designed by each PEPOP method.

#### Probability

If X is a surface-accessible aa (alanine, cysteine, …, tyrosine), n_X_ and p_X_ represent the number of occurrences of X in the protein and the peptide, respectively. The probability to obtain a specific peptide sequence by chance is thus given by the following formula:

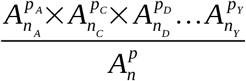

with:

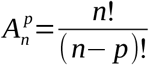

where *n*=*n_A_* + *n_C_* + *n_P_* +…+*n_Y_* is the number of surface-accessible aa in the protein and *p*= *p_A_* + *p_C_* + *p_P_* +…+ *p_Y_* the number of aa in the peptide.

### SUPERFICIAL

The aim of SUPERFICIAL [36] is to design peptides that mimic regions at the surface of a given protein, starting from its 3D structure. SUPERFICIAL first computes the surface-accessibility of each aa and then builds segments as surface-accessible and contiguous aa sequences. Peptides can be made of several segments close in space, linked together in order to conserve the local conformation of the targeted protein surface. SUPERFICIAL finds the linkers by calculating the number (not the type) of aa needed to link two segments, based on the distances and angles between their C- and N-termini.

## Acknowledgements

We warmly thank Dr Ponomarenko for providing the dataset of antigen-antibody complexes. We thank Dr P. Lapalud for her collaboration, D. Jean for his precious technical contribution and Y. Crouineau for his relevant advices in graphiscs. We thank Dr E. Dupas for her contribution. This work was supported by grants from the Ministry of Research (France); University Paris Diderot, Sorbonne Paris Cité (France); the National Institute for Blood Transfusion (INTS, France); the Institute for Health and Medical Research (INSERM, France); and a Labex GR-Ex grant (France) to Alexandre de Brevern. The Labex GR-Ex, reference ANR-11-LABX-0051, is funded by the program “Investissements d’avenir” of the French National Research Agency, reference ANR-11-IDEX-0005-02.

